# The influence of insulin therapy on measurements of optical coherence tomography parameters in diabetic patients without retinopathy

**DOI:** 10.1101/2020.01.31.928184

**Authors:** Deyuan Zhou, Wei Wang, Rouxi Zhou, Miao He, Xia Gong, Yuting Li, Wenyong Huang

## Abstract

**Purpose:** To determine whether there was a change in the fundus of the eye in diabetic patients without retinopathy after insulin therapy.

**Methods:** The diabetic patients without retinopathy were included in this study. A swept-source optical coherence tomography/angiography (SS-OCT/A) was used to obtain the measurements of macular retinal nerve fibre layer (mRNFL) thickness, ganglion cell-inner plexiform layer (GC-IPL) thickness, retinal thickness (RT), macular choroidal thickness (MCT), peripapillary retinal nerve fibre layer (pRNFL) thickness, peripapillary choroidal thickness (PCT), and perfused vascular density (PVD). Univariable and multivariable regression analyses were performed to explore the influence of insulin use on measurements of OCT/A.

**Results:** A total of 1140 patients used insulin (using group), and 126 patients did not use insulin (without group). The average MCT of the using group was 171.3±67.8 μm, which was thinner than that in the without group (190.2±74.7 μm) (P=0.012). The average PVD of the using group (48.0±2.1 μm) was less than that in the without group (48.7±2.1 μm) (P<0.001). After adjusting for age, gender, axial length, duration, HbA1c, systolic blood pressure, diastolic blood pressure, cholesterol, serum creatinine, insulin use was significantly associated with thinner MCT (beta=-16.12μm; 95%CI:-29.42, −2.81μm; P= 0.018) and lower PVD (beta=-0.79; 95%CI: −1.22, −0.36; P<0.001).

**Conclusion:** The use of insulin by diabetic patients without retinopathy might decrease the MCT and PVD compared to patients who did not use insulin, which helps to better understanding the role of insulin use on higher risk for diabetic retinopathy.

## Introduction

Type 2 diabetes mellitus (T2DM) poses a major public challenge, which is one of the most common chronic diseases affecting nearly half a billion people worldwide.^1^ Diabetic retinopathy (DR) is the most common ocular complication of T2DM and the leading cause of blindness in working-age population.^2^ Diabetic macular oedema (DME) is the main cause of vision loss of DR.^3^ There were 92.6 million people with DR and 20.6 million people with DME globally.^4^ The primary goal of DR management is to prevent its occurrence, thus it is crucial to identify risk factors and explore pathogenesis of DR.^3^ Though development of DR is multifactorial, hyper-glycemia was considered to be the most important risk factor for DR and DME.

Insulin is one of the most reliable therapies for diabetes, which has been proven to maintain a normal blood glucose concentration and protect the insular function, therefore, significantly reduce the risk of the development or the progression of DR.^5^ Unfortunately, some recent studies reported that insulin use was associated with high risk of hypertension, limb ischemia, and various carcinomas.^6, 7^ There were also studies suggested that insulin use exacerbates DR or induces early retinal deterioration that could lead to DR in a diabetic patient.^8–10^ A meta-analysis of 19107 subjects demonstrated that insulin use was an independent risk factor for DR (relative risk = 2.30).^8^ Another meta-analysis involving 202 905 participants found that insulin use was associated with 2.42 times higher odds for DME.^10^ To determine the mechanism by which insulin affects the retina, determination of any changes to the physiological characteristics of the retina after insulin injections is required. However, little information was available for this topic and the effect of insulin use on retinal structure and function needs advanced research.^11, 12^

The introduction of optical coherence tomography (OCT) enable visualization and segmentation of each layer of retina and choroid, which has become an essential tool in management of DR patients. The OCT angiography (OCTA) provides the quantitative and qualitative analyses of retinal microvasculature without dyes injection.^13^ A variety of studies have reported that changes of inner retinal layers, choroid thinning, and inadequate perfusion of retinal vessels play a key role in onset and progression of DR.^14, 15^ However, the influence of insulin use on OCT and OCTA parameters have not explored in T2DM without DR. Therefore, the objective of this was to analyse the OCT parameters in a large sample of T2DM patients without DR based on insulin use and to investigate the difference in these parameters between patients who used insulin and patients who did not use insulin.

## Methods

### Participants

This cross-sectional study was performed at the Zhongshan Ophthalmic Centre (ZOC) of Sun Yat-sen University in China. The protocol of the study was approved by the ZOC Institute Ethics Committee. The study was performed according to the tenets of the Declaration of Helsinki. All participants signed written informed consent forms before enrolment. Patients with T2DM in the local government registry database were invited to participate in this study. Eligible patients were those with T2DM aged 40 to 80 years old who had no previous eye treatment or surgery. Patients were excluded if they had any of the following: a spherical equivalent (SE) of −6.0 D or more or a cylinder degree of 3.0 D or more in any eye; prior retinal surgery, intravitreal injection, or laser photocoagulation; a history of cataract surgery or other intraocular surgery; ocular pathology that interferes with imaging (e.g., dense cataracts or corneal ulcers); eyes with ungradable OCTA images, structural OCT images, or colour fundus photographs; retinal diseases or neuropathy other than DR (e.g., epiretinal membrane, glaucoma, retinal vein occlusion, or wet age-related macular degeneration).

### General information and laboratory tests

General information was collected via standardized questionnaires, including age, sex, duration of diabetes, medication adherence, comorbidities, and lifestyle factors. The duration of diabetes, history of associated comorbidities, and other diabetic complications were further confirmed by the information from each patient’s medical record. A history of cardiovascular disease (CVD) was defined as a history of coronary heart disease or stroke. Height, weight, systolic blood pressure (SBP), and diastolic blood pressure (DBP) were measured by standardized protocol. Blood and urine samples were obtained from all patients for testing the following parameters: haemoglobin A1c (HbA1c), total cholesterol (TC), high-density lipoprotein cholesterol (HDLC), low-density lipoprotein cholesterol (LDLC), triglycerides (TG), serum creatinine, C-reactive protein (CRP), and urine microalbuminuria.

### Ocular examination

All participants underwent a comprehensive ophthalmic examination, including best corrected visual acuity (BCVA), intraocular pressure, refractive error, ocular biometrical measurements, slit-lamp biomicroscopy, and a dilated fundus examination. The BCVA was converted to the logarithm of the minimum angle of resolution (LogMAR) for statistical analysis. The dilated fundus photography was performed by using standard 7 field protocol defined by Early Treatment Diabetic Retinopathy Study (ETDRS). The DR severity and DME was assessed by retinal specialists according to the International Clinical Diabetic Retinopathy Disease Severity Scale and the Diabetic Macular Edema Disease Severity Scale.^16^ DR severity was categorised as absent, mild, moderate, or severe non-proliferative DR or proliferative DR, and DME was classified as absent or present. The ocular biometrical measurements were obtained using optical low-coherence reflectometry (Lenstar LS900; Haag-Streit AG, Koeniz, Switzerland), including central corneal thickness (CCT), anterior chamber depth (ACD), lens thickness (LT), axial length (AL), and corneal diameter (CD). The refractive error was measured by an autorefractor (KR8800; Topcon, Tokyo, Japan) after bilateral pupil dilation.

### OCT and OCTA imaging

Bothe OCT and OCTA imaging were performed by a commercial swept-source OCT (SS-OCT) device (DRI OCT-2 Triton; Topcon, Tokyo, Japan) to obtain high-definition retina and choroid structure images and vessel OCTA maps. The device had a speed of 100,000 A-scans/s and yielded an 8-μm axial resolution in tissue. All OCT scans were performed by the same experienced technician who was blind to the study protocol. Before the scan was conducted, we verified that none of the patients had consumed caffeine or alcohol or had taken analgesic medications for at least 24 h prior to the procedure. The method of structure imaging with SS-OCT and vascular imaging with SS-OCTA have been described in detail elsewhere.^17, 18^

Three-dimensional (3-D) imaging scans were obtained using the 7×7 mm raster scan protocol centred on the macula. The resultant images were analysed by the automated layer segmentation software (version 9.12.003.04) that was built into the SS-OCT system. The macular retinal nerve fibre layer (mRNFL) thickness, ganglion cell-inner plexiform layer (GC-IPL) thickness, retinal thickness (RT), macular choroidal thickness (MCT) automatically calculated and displayed in the nine subfields defined by the ETDRS (Figure 1). The average mRNFL thickness, GC-IPL thickness, RT, and MCT thickness in all nine grids were calculated for statistical analyses. The peripapillary imaging scans were obtained using 3.4 mm circle scan protocol centred on optic nerve head. The average peripapillary retinal nerve fibre layer (pRNFL) thickness and peripapillary choroidal thickness (PCT) were obtained for statistical analyses. Only patients with eligible images (i.e., image quality >50 without eye movement, artefacts, or segmentation failure) for both eyes were included in the study.

**Figure 1.**
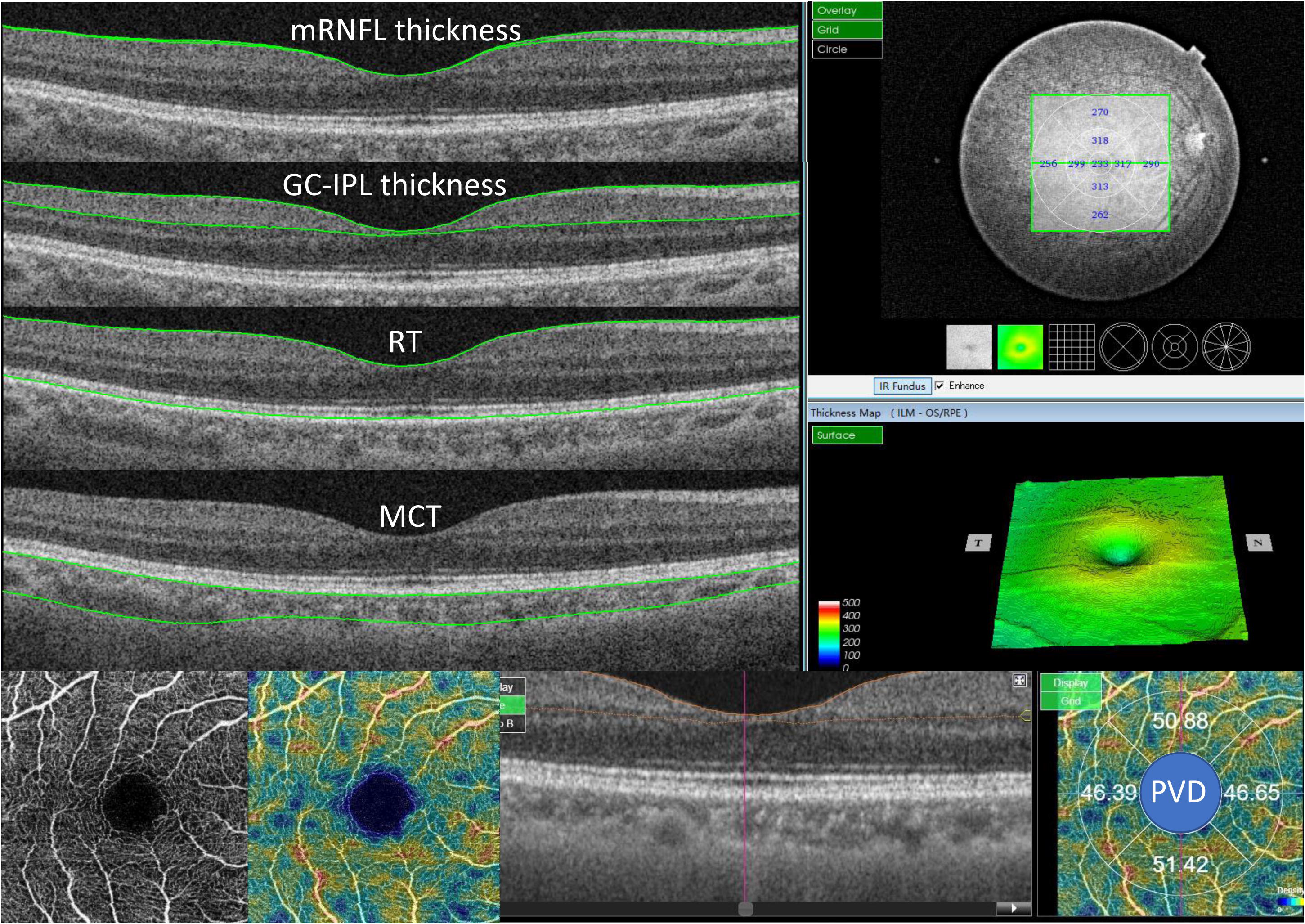
Diagrams of measurements of the swept-source optical coherence tomography / angiography (SS-OCT/A) in grids of Early Treatment Diabetic Retinopathy Study. mRNFL =macular retinal nerve fibre layer; GC-IPL=ganglion cell-inner plexiform layer; RT=retinal thickness; MCT=macular choroidal thickness; PCT= peripapillary choroidal thickness; pRNFL= peripapillary retinal nerve fibre layer; PVD=parafoveal vessel density.

Volumetric OCTA scans centered on the fovea were obtained with a scan area of 3×3 mm, which contained 320×320 A-scans. The latest built-in software (IMAGEnet6) was used to generate OCT-angiograms, which can improve the detection sensitivity of low blood flow and reduce motion artefacts without compromising axial resolution. Parafoveal vessel density (PVD) were calculated based on the ETDRS grid. The parafoveal region was defined as an annulus with an outer diameter of 3 mm and an inner diameter of 1 mm (Figure 1). Vessel density was calculated as the percentage of the area not defined as non-perfusion regions over the total area within the region of interest. An image quality score ranging from 0 to 100 was provided by the software for each volumetric OCT scan. Images with quality score > 50, without artefacts and slab errors were included in the study.

### Statistical analyses

The Kolmogorov-Smirnov test was performed to verify a normal distribution. The student t test and chi-square test were used to compare the characteristics and OCT parameters between patients using insulin and patients not using insulin (companion non-parametric tests were considered if necessary). Three linear regression models were used to analyse the association of OCT parameters and insulin use. Model 1 was a univariable linear regression to assess the associations of OCT parameters with insulin use. Our previous studies found that OCT parameters are associated with age, sex, AL, and other factors.^19, 20^ Therefore, model 2 was adjusted for age and sex, and model 3 was further adjusted for other confounding factors, including AL, duration of diabetes, HbA1c, SBP, DBP, total cholesterol, and serum creatinine, in the multivariable analyses. All analyses were performed using Stata Version 14.0 (Stata Corporation, College Station, TX, USA). A P value of <0.05 was considered statistically significant.

## Results

Table 1 shows the demographic and clinical characteristics between the insulin using group and the without group. The insulin using group had an average T2DM duration of 7.5±6.2 years, which was shorter than that in the without group (12.0±6.7 years) (P<0.001). The HbA1c level of the insulin using group (6.7±1.2%) was lower than that in the without group (7.4±1.6%) (P<0.001). The blood pressure was significantly lower in insulin using group than that in the without group, with SBP (133.2±18.5 mmHg versus 138.7 ±19.7 mmHg; P=0.002) and DBP (70.6±10.1 mmHg versus 73.0±11.2 mmHg; P=0.012). The BCVA of the insulin using group was 0.2±0.1 LogMAR unit, which was lower than that of the without group (0.3±0.2) (P=0.048). The SE value was 0.2±2.2 in the insulin using group compared to −0.5 ±2.5 in the without group (P=0.004). Both groups had similar BMI, AL, ACD, and other clinical parameters (Table 1).

**Table 1.** Demographic and clinical characteristics of type 2 diabetic patients without retinopathy.

| Characteristics | All | Use insulin | No insulin | P-value |
| --- | --- | --- | --- | --- |
| No. of subjects | 1266 | 1140(90.1%) | 126(9.9%) | - |
| Gender (%) |  |  |  | 0.263 |
| Female | 762(60.19%) | 692(60.7%) | 70(55.56%) |  |
| Male | 504(39.81%) | 448(39.3%) | 56(44.44%) |  |
| Age, year | 64.6±7.9 | 64.7±7.8 | 63.4±8.5 | 0.080 |
| Duration of diabetes, year | 7.9±6.4 | 7.5±6.2 | 12.0±6.7 | **<0.001** |
| HbA1c, % | 6.8±1.3 | 6.7±1.2 | 7.4±1.6 | **<0.001** |
| BMI, kg/m<sup>2</sup> | 24.7±3.3 | 24.8±3.2 | 24.6±3.8 | 0.565 |
| Systolic BP, mm Hg | 133.8±18.6 | 133.2±18.5 | 138.7±19.7 | **0.002** |
| Diastolic BP, mm Hg | 70.9±10.2 | 70.6±10.1 | 73.0±11.2 | **0.012** |
| Total cholesterol, mmol/L | 4.8±1.0 | 4.8±1.0 | 4.8±1.0 | 0.814 |
| Serum creatinine, µmol/L | 70.9±19.5 | 71.0±19.2 | 70.0±21.8 | 0.605 |
| HDL-c, mmol/L | 1.3±0.4 | 1.3±0.4 | 1.3±0.4 | 0.964 |
| LDL-c, mmol/L | 3.1±0.9 | 3.1±0.9 | 3.1±0.9 | 0.628 |
| Triglycerides, mmol/L | 2.3±1.6 | 2.4±1.6 | 2.1±1.4 | 0.122 |
| Uric acid, µmol/L | 369.5±95.6 | 370.9±95.9 | 356.6±91.7 | 0.113 |
| CRP, mg/L | 2.9±7.6 | 2.9±7.8 | 2.9±5.1 | 0.962 |
| MAU, mg/mL | 3.8±12.8 | 3.9±13.2 | 3.5±9.0 | 0.723 |
| BCVA, logMAR | 0.2±0.1 | 0.2±0.1 | 0.3±0.2 | **0.048** |
| IOP, mmHg | 16.2±2.8 | 16.2±2.8 | 15.8±2.5 | 0.181 |
| SE, diopter | 0.1±2.3 | 0.2±2.2 | -0.5±2.5 | **0.004** |
| CCT, µm | 546.6±31.6 | 546.6±31.9 | 546.5±29.1 | 0.963 |
| AL, mm | 11.6±0.4 | 11.6±0.4 | 11.7±0.4 | 0.211 |
| CD, mm | 44.2±1.5 | 44.2±1.5 | 44.2±1.5 | 0.915 |
| ACD, mm | 2.4±0.4 | 2.4±0.4 | 2.5±0.5 | 0.056 |

Data are expressed as the mean ± SD or %.
BMI=body mass index; BP = blood pressure; HDL-c= high density lipoprotein cholesterol; LDL-c= low density lipoprotein cholesterol; CRP=C-reactive protein; MAU= microalbuminuria; BCVA=best corrected visual acuity; IOP=intraocular pressure; SE=spherical equivalent; CCT= central corneal thickness; AL= axial length; CD=corneal diameter; ACD=anterior chamber depth;.
Bold indicates statistical significance.

Table 2 presents the differences of OCT parameters between insulin using group and without using group. Both groups had similar values of average mRNFL thickness, macular GC-IPL thickness, RT, PCT, and pRNFL thickness. Patients in the insulin using group had a significantly lower average MCT than that in the without group (171.3±67.8 μm versus 190.2 ±74.7 μm; P<0.012). Furthermore, the mean PVD of the using group was significantly lower than that in the without group (48.0%±2.1% versus 48.7%±2.1%; P<0.001).

**Table 2.** Differences of average measurements of OCT and OCTA parameters between the insulin using group and without group.

| **Parameters** | **No insulin** | **Use insulin** | **P-value** |
| --- | --- | --- | --- |
| Average mRNFL thickness, $\mu\text{m}$ | 28.4 ±4.0 | 29.1 ±4.3 | 0.081 |
| Average GC-IPL thickness, $\mu\text{m}$ | 72.4 ±7.0 | 72.2 ±7.0 | 0.824 |
| Average RT, $\mu\text{m}$ | 273.7 ±19.0 | 273.5 ±20.0 | 0.923 |
| Average MCT, $\mu\text{m}$ | 190.2 ±74.7 | 171.3 ±67.8 | **0.012** |
| Average PCT, $\mu\text{m}$ | 114.5 ±54.1 | 104.7 ±54.1 | 0.076 |
| Average pRNFL thickness, $\mu\text{m}$ | 110.7 ±13.5 | 109.3 ±11.6 | 0.285 |
| Average PVD, % | 48.7 ±2.1 | 48.0 ±2.1 | **<0.001** |

mRNFL =macular retinal nerve fibre layer; GC-IPL=ganglion cell-inner plexiform layer; RT=retinal thickness; MCT=macular choroidal thickness; PCT= peripapillary choroidal thickness; pRNFL= peripapillary retinal nerve fibre layer; PVD=parafoveal vessel density.
Bold indicates statistical significance.

Table 3 summaries the influence of insulin using on OCT parameters by using regression analyses. The insulin using was associated with a thinner MCT of −18.89μm (95%CI: −33.56, −4.22; P=0.012) and a lower PVD of −0.69% (95%CI: −1.09%, −0.29%, P=0.001). After adjusting for age and sex, the results were similar. Further adjusting for AL, disease duration, HbA1c, SBP, DBP, total cholesterol, and serum creatinine, the results confirmed that insulin using was independently related to a reduced MCT by −16.12μm (95%CI: −29.42, −2.81; P=0.018) and PVD by −0.79% (95%CI:-1.22, −0.36; P<0.001).

**Table 3.** Univariable and multivariable regression analyses exploring the association between insulin use and measurements by optical coherence tomography.

| Parameters | $\beta$ | 95%CI | | P-value |
| --- | --- | --- | --- | --- |
|  |  | Lower | Upper |  |
| **Model 1\*** |  |  |  |  |
| Average mRNFL thickness, $\mu\text{m}$ | 0.72 | -0.09 | 1.53 | 0.081 |
| Average GC-IPL thickness, $\mu\text{m}$ | -0.16 | -1.55 | 1.24 | 0.824 |
| Average retinal thickness, $\mu\text{m}$ | -0.19 | -3.98 | 3.61 | 0.923 |
| Average MCT, $\mu\text{m}$ | -18.89 | -33.56 | -4.22 | **0.012** |
| Average PCT, $\mu\text{m}$ | -9.79 | -20.60 | 1.01 | 0.076 |
| Average pRNFL thickness, $\mu\text{m}$ | -1.45 | -4.11 | 1.21 | 0.285 |
| Average PVD, % | -0.69 | -1.09 | -0.29 | **0.001** |
| **Model 2†** |  |  |  |  |
| Average mRNFL thickness, $\mu\text{m}$ | 0.66 | -0.14 | 1.46 | 0.104 |
| Average GC-IPL thickness, $\mu\text{m}$ | -0.37 | -1.71 | 0.96 | 0.583 |
| Average retinal thickness, $\mu\text{m}$ | -1.16 | -4.74 | 2.42 | 0.526 |
| Average MCT, $\mu\text{m}$ | -21.98 | -35.72 | -8.24 | **0.002** |
| Average PCT, $\mu\text{m}$ | -12.11 | -22.29 | -1.92 | 0.02 |
| Average pRNFL thickness, $\mu\text{m}$ | -1.68 | -4.30 | 0.93 | 0.207 |
| Average PVD, % | -0.71 | -1.11 | -0.31 | **<0.001** |
| **Model 3‡** |  |  |  |  |
| Average mRNFL thickness, $\mu\text{m}$ | 0.36 | -0.47 | 1.20 | 0.393 |
| Average GC-IPL thickness, $\mu\text{m}$ | 0.10 | -1.31 | 1.51 | 0.891 |
| Average retinal thickness, $\mu\text{m}$ | -0.08 | -3.92 | 3.75 | 0.966 |
| Average MCT, $\mu\text{m}$ | -16.12 | -29.42 | -2.81 | **0.018** |
| Average PCT, $\mu\text{m}$ | -7.57 | -18.35 | 3.21 | 0.168 |
| Average pRNFL thickness, $\mu\text{m}$ | -0.14 | -2.87 | 2.58 | 0.918 |
| Average PVD, % | -0.79 | -1.22 | -0.36 | **<0.001** |

mRNFL =macular retinal nerve fibre layer; GC-IPL=ganglion cell-inner plexiform layer; RT=retinal thickness; MCT=macular choroidal thickness; PCT= peripapillary choroidal thickness; pRNFL= peripapillary retinal nerve fibre layer; PVD=parafoveal vessel density.
Model 1: univariable linear regression
Model 2: adjusted for age and sex.
Model 3: adjusted for age, gender, axial length, duration, HbA1c, systolic blood pressure, diastolic blood pressure, cholesterol, serum creatinine.
Bold indicates statistical significance.

## Discussion

DR remains a major cause of irreversible blindness in the working age population. Though the strictly controlling blood glucose delays the development and progression of DR in the long term, it was reported that insulin use led to blurred vision initially and increased DR and DME risk in patients without retinopathy.^8, 10, 21^ The diabetes control and complications trial (DCCT) found that insulin use led to transient DR worsen in the short-term, which was more observed in intensive insulin therapy group than that in regular insulin therapy group.^22^ The advance of OCT techniques enables to visualize cross-sectional retina and retinal microvasculature in vivo. Thus, it is important to know if there are any differences in these two parameters and what these differences are, which may lead to a method for determining the basic mechanism of how insulin affects the fundus of the eye. In this study, we discovered that the use of insulin by diabetic patients without retinopathy alters some of the fundus OCT parameters compared to diabetic patients who did not use insulin. As far as we known, this is the first report to evaluate the influence of insulin use on OCT and OCTA parameters in T2DM without DR.

We found that the insulin use was related to a thinning choroid. The choroid is a vascular tissue of the richest blood flow in the eyeball, which has implicated in a variety retinal and choroidal diseases such as DR, macular degeneration, central serous chorioretinopathy.^23^ In this study, patients who used insulin had a lower MCT than patients who did not use insulin. This reduced MCT might contribute to the relationship between insulin use and DR/DME risk. A number of studies have reported that the MCT was reduced significantly in DR and correlated with severity of DR and DME.^18, 24–26^ Several factors may contribute to the relationship between insulin use and MCT thinning. Sheng et al.^27^ demonstrated that insulin use prevented the choroidal thickening caused by wearing positive lenses in chicken eyes by upregulating expression of diffusible factors, including vascular endothelial growth factor (VEGF), insulin growth factor 1 (IGF-1), and IGF-2. Hernandez et al.^21^ reported that insulin therapy resulted in increased macular volume through the reduction of serum soluble receptor of VEGF (Flt-1, VEGFR-1). Furthermore, experimental study in diabetic mice found that insulin increased microvascular permeability by activating nitric oxide, HIF-1α, and epidermal growth factor (EGF) receptor signal pathways.^28,29^ Zhang et al.^30^ reported that serum IGF-1 was independently related to MCT. Other studies reported that MCT was significantly influenced by levels of aqueous and serum VEGF.^31, 32^ Further studies are needed to unravel the underling mechanism.

We also discovered that the PVD of the using group was lower than that of the without group, although it was a minimal difference. Previous studies reported a decrease in pulsatile ocular blood flow was observed in the early stage of observable DR, and a lower PVD was related to the development and progression of DR.^33–35^ A prior study reported that insulin could induce vascular endothelial production of nitric oxide, which leads to capillary recruitment, vasodilation, and increased blood flow.^36^ However, our study demonstrated that insulin use could decrease the ocular PVD. Therefore, a reduction in PVD might be caused by other mechanism. We assume that insulin could activate apamin-sensitive Ca^2+^-activated potassium channels, and then hyperpolarization in retinal capillary pericytes could cause angiectasis and reduce blood flow to the retina, as shown in a previous study.^37^ In addition, the expression of VEGF may play a role. It was reported that aqueous level of VEGF in DR was significantly associated with capillary nonperfusion area measured by OCTA.^38^ The mechanism by which insulin use decreased retina blood flow needs further exploration in future in vitro and vivo studies.

The findings had clinical and research implications. This study showed a lower MCT and lower PVD in diabetic patients without retinopathy who used insulin than those of patients who did not use insulin, which revealed the effect of insulin on the ocular fundus. This finding was consistent with previous studies reporting higher risk for DR and DME after insulin therapy, which enables a better understanding of the role of insulin use in development of DR and DME.^8–10^ As the participants had a long history of diabetes and insulin therapy, we can better understand the effects of continuous insulin therapy on the eye. OCTA is a non-invasive imaging modality that enables to detect the blood flow without dye injection. Since the introduction of OCTA, a number of studies have evaluated the PVD changes in diabetic patients.^39, 40^ Various factors such as age, sex, axial length, serum creatine have been reported to influence OCTA parameters. However, few studies have considered the effect of insulin when studying DR.^39, 40^ This study suggested that future OCTA studies should adjusted for insulin use in the statistical analyses.

This study had some limitations. Firstly, the participants were all T2DM patients, and the results we discovered in this study might not apply to patients with type 1 diabetes mellitus. Secondly, we only observed the difference between the two groups at one point in time. Other changes may exist during the times we did not observe. However, we assume that given the large sample size and adjusting various confounding factors, the conclusions are robust and reliable. Thirdly, the disease duration of the using group was shorter than that of the without group. Therefore, blood glucose control might have been worse in the without group, which might have had negative effects on the eyes and affected our study. Finally, because the nature of cross-sectional design, a causal relationship between insulin use and OCTA changes could not be established. Additional prospective cohort studies are still warranted to confirm our results.

In summary, we provide evidence that insulin use was associated with a decrease in MCT and PVD in T2DM patients without DR, which enables a better understanding of the role of insulin use in the development of DR and DME. The mechanism by which insulin use decreased choroid and retina blood flow needs further exploration in future in vitro and vivo studies. Additional large-scale longitudinal studies are needed to confirm our findings.

## Acknowledgments

This study was supported by the National Natural Science Foundation of China (81570843; 81530028; 81721003). The funding organizations had no role in the design or conduct of the study; collection, management, analysis, and interpretation of the data; preparation, review, or approval of the manuscript; and decision to submit the manuscript for publication.

## Authors’ Contributions

WW and WH had full access to all the data in the study and take responsibility for the integrity of the data and the accuracy of the data analysis. Study concept and design: WW, MH, WH. Acquisition, analysis, or interpretation of data: DZ, YT, XG, RZ, WH. Drafting of the manuscript: DZ, WW. Critical revision of the manuscript for important intellectual content: All authors. Statistical analysis: WW. Obtained funding: WH. Administrative, technical, or material support: MH, WW. Study supervision: MH.

